# Evolutionary and methodological considerations when interpreting gene presence-absence variation in pangenomes

**DOI:** 10.1101/2025.08.14.670405

**Authors:** Tomáš Brůna, Avinash Sreedasyam, Avril M. Harder, John T. Lovell

**Affiliations:** U.S. Department of Energy Joint Genome Institute, Lawrence Berkeley National Laboratory, Berkeley, CA 94720, USA; Genome Sequencing Center, HudsonAlpha Institute for Biotechnology, Huntsville, AL 35806, USA

## Abstract

While graph-based pangenomes have become a standard and interoperable foundation for comparisons across multiple reference genomes, integrating protein-coding gene annotations across pangenomes in a single ‘pangene set’ remains challenging, both because of methodological inconsistency and biological presence-absence variation (PAV). Here, we review and experimentally evaluate the root of genome annotation and pangene set inconsistency using two polyploid plant pangenomes: cotton and soybean, which were chosen because of their existing diverse high-quality genomic resources and the known importance of gene presence-absence variation in their respective breeding programs. We first demonstrate that building pangene sets across different genome resources is highly error prone: PAV calculated directly from the genome annotations hosted on public repositories recapitulates structure in annotation methods and not biological sequence differences. Re-annotation of all genomes with a single identical pipeline largely resolves the broadest stroke issues; however, substantial challenges remain, including a surprisingly common case where exactly identical sequences have different gene model structural annotations. Combined, these results clearly show that pangenome gene model annotations must be carefully integrated before any biological inference can be made regarding sequence evolution, gene copy-number, or presence-absence variation.

## Introduction

Pan-genome resources, or the representation of all available DNA sequences across groups of organisms in a single catalog (hereon ‘pangenomes’), are rapidly becoming the standard foundation for genetics research in medicine, agriculture, biotechnology, ecology, and evolution (Jayakodi et al. 2024). This paradigm shift has been made possible by advances in long-read sequencing technology and assembly algorithms. Multiple functionally complete genomes are now available for many species. The tools to construct, compare and visualize pangenomes have also rapidly advanced in a handful of heavily invested model systems. In these cases ‘graph’ representations have been shown to improve variant detection (Zhou et al. 2022; Wang et al. 2022), providing clues to evolutionary processes and sequence diversity in previously inaccessible regions of the genome. In contrast to the technological advances that aid in the construction and integration of multiple nearly complete and consistent genome assemblies, *de novo* annotation of putatively functional protein coding gene models remains challenging and expensive (Vuruputoor et al. 2023; Abbas et al. 2024).

Typically, protein-coding gene annotations are guided by one of three main data types: (1) ‘homology’ similarity to well-supported but phylogenetically distant protein sequences, (2) ‘*ab initio*’ gene structure evidence, or (3) short-read and full-length RNA sequencing-based ‘transcriptomes’ (Brůna et al. 2024). However, each of these evidence structures can produce high error rates: (1) homology-driven methods are unlikely to discover true new genes or gain-of-function variants (high false negative), (2) *ab initio*-driven methods tend to have very high rates of false-positive ‘artifact’ genes, and (3) transcriptome-driven methods often produce variable results depending on the amount, type, and source (but see below) of sequencing support.

Recently, Gabriel et al. (2024) demonstrated that ‘integrative’ annotation methods, which expend the substantial compute and sequencing costs to combine the three main evidence supports (homology, *ab initio*, and transcriptome), are likely to best discover true gene presences and minimize false positives. However, given the costs of integrative pipelines, many annotations of pangenomes now heavily rely on *ab initio* and/or homology support (e.g., Helixer (Holst et al. 2023) or Galba (Brůna et al. 2023)). In the extreme case, some pangene sets are generated from gene model propagation ‘lift-over’ from a single reference to all members of the pangenome (e.g., LiftOff (Shumate and Salzberg 2021)). However, annotation methods that rely solely on lifting reference gene models are incapable of new gene discovery and are therefore not appropriate for any but the most homogenous pangenomes, which are probably best represented by a single reference genome.

Individual gene annotation quality is typically assessed through complementary metrics: completeness measures such as BUSCO (Manni et al. 2021), which quantify the presence of expected conserved orthologs as true-positives, and precision measures such as PSAURON (Sommer et al. 2025), which estimate false-discovery rates by identifying spurious predictions. Because genomic inquiries in many systems are beginning to rely on multiple references, consistency of gene annotations for orthologous sequences across references is a third and particularly important quality metric. Such consistency is critical for evolutionary inference and practical applications of pangenomes: often the most important product from a pangenome is functional sequence variation across a set of genes (i.e., the ‘pangene set’) that share an evolutionary common ancestor (i.e., pairwise ‘orthologs’ or sets of genes in ‘orthogroups’).

When approaching pangenome annotations, these three aspects — completeness, precision, and consistency — must be considered and balanced. The most consistent annotations (e.g., lift-overs) will by definition be unable to find new genes and thus suffer from false-negatives in gene lists. Conversely, highly sensitive approaches that identify all possible coding sequences typically generate high false discovery rates in individual genomes and correspondingly low consistency. However, the extent to which pangene sets are consistent, both across annotation methods and within pangenomes annotated by identical approaches, is not well understood.

In this study, we investigate the extent to which gene presence-absence variation (PAV) in pangenomes is driven by biological versus methodological factors, with particular attention to the challenges of post hoc integration of gene annotations generated with diverse computational methods and input resources. We first examine PAV patterns across taxonomic groups with different evolutionary histories, comparing plants and animals at various divergence timescales. We then analyze publicly available cotton and soybean pangenomes to quantify how annotation methodology impacts inferences of relatedness and gene content variation. Our results demonstrate that while biological factors certainly contribute to gene PAV, methodological inconsistencies can dramatically inflate estimates of pangenome “openness” and potentially mislead evolutionary and functional interpretations. These findings have important implications for pangenome construction, annotation strategies, and downstream applications in breeding, comparative genomics, and evolutionary biology.

## Results and Discussion

### What is the scale of gene presence-absence variation across annotations?

The relative degree to which two genome annotations are ‘consistent’ depends largely on two factors: (1) the *biological* predisposition of genomes to lose or gain true protein-coding genes, and (2) the *technical* ability for annotation method to identically call similarly structured genes models from orthologous DNA sequences. Biological causes of gene presence-absence variation abound in the literature (e.g., Li and Lee (2023); Hurgobin et al. (2018)), and it has been hypothesized that lineages that have undergone whole-genome duplications (‘WGDs’), like most plants, fungi (Albertin and Marullo 2012), insects (Li et al. 2018), but not most vertebrates (see Glasauer and Neuhauss (2014)), will harbor biologically relevant gene copy number (CNV) and presence-absence (PAV) variation that may drive speciation and complex trait evolution (Lye and Purugganan 2019).

We tested the extent that gene PAV and CNV vary across two taxonomic groups (flowering plants, animals) and four levels of divergence times (within species, 6-7My, ∼100My, > 150My; Table 1). Unsurprisingly, within both flowering plants and amniotes, gene PAV becomes increasingly common with increasing divergence time—more diverged genomes share fewer orthologs, likely due to sequence deletions, pseudogenization and other forms of large-scale mutation followed by selection or drift. However, the absolute amount of gene PAV and the degree that PAV is affected by divergence time was remarkably different between plants and animals. For example, the most closely related plant contrast (between two maize genotypes) exhibited similar PAV to the ∼90My diverged human and mouse genomes. While it is possible that some of this variation among the maize genomes could be due to their ∼10My WGD, the ∼7My diverged tobacco species that lack a recent WGD had a higher proportion of PAV gene families than the 300My diverged human-chicken contrast. Consistent with previous observations (Roeder et al. 2025), fewer than half of all genes were found in 1:1 ortholog families between the two major flowering plant lineages.

**Table 1.** Copy number and presence-absence variation between pairs of flowering plant and vertebrate genomes. While not exactly the same, each contrast is between pairs of genomes that have diverged by approximately the same number of years ^+^as estimated by timetree.org. Contrasts are as follows: *Canus lupus familiaris* (domesticated dog) ‘Tasha’ vs. ‘Ros’; ^2^*Zea mays* (Maize) ‘B73’ vs. ‘B97’; ^3^*Homo sapiens* (human) ‘CHM-13’ vs. Pan troglodytes (Chimpanzee) ‘AG18354’; ^4^*Nicotiana sylvestris* vs *Nicotiana tomentosiformis* (wild tobacco species); ^5^*Homo sapiens* vs. Mus muscus (house mouse) ‘GRCm39’; ^6^*Nicotiana sylvestris* vs *Coffea arabica* (coffee); ^7^*Homo sapiens* vs *Gallus gallus* (chicken) ‘bGalGal1’; ^8^*Oryza sativa* (rice) vs. *Coffea arabica*. Gene PAV is defined with phylogenetically hierarchical orthogroups from OrthoFinder. All contrasts are between NCBI ‘refseq’ genomes that were both annotated by the NCBI pipeline *except Maize (from maizegdb), which were both annotated by a separate method that was applied identically to both B73 and B97, since no plant species has two NCBI-refseq annotations. The last line contains the % PAV genes, which is colored by distance away from the mean value across all comparisons (increasing blue: < mean, increasing red: > mean).

|  | Dogs<sup>1</sup> | Maize<sup>2</sup> | Primates<sup>3</sup> | Tobacco<sup>4</sup> | Placentals<sup>5</sup> | Malvids<sup>6</sup> | Amniotes<sup>7</sup> | Angiosperms<sup>8</sup> |
| --- | --- | --- | --- | --- | --- | --- | --- | --- |
| Est. Divergence | <1Mya |  | 6-7Mya |  | ~90-100Mya |  | ~300M, ~160Mya |  |
| *n* 1:1 orthologs | 18,674 | 23,178 | 17,119 | 19,452 | 15,596 | 12,204 | 11,686 | 2,143 |
| *n* CNV orthogroups | 742 | 19,837 | 1,378 | 4,860 | 1,383 | 2,256 | 1,870 | 8.852 |
| *n* PAV orthogroups | 768 | 10,434 | 1,962 | 7,959 | 2,484 | 6,975 | 4,262 | 16,053 |
| % 1:1 genes | 90.3% | 32.1% | 79.0% | 44.3% | 71.6% | 50.1 | 60.3% | 5.4% |
| % PAV genes | 3.2% | 11.6% | 8.4% | 23.3% | 14.3% | 30.4% | 21.3% | 53.5% |

Like among these biological contrasts, in some cases, the technical causes of gene PAV is potentially obvious. For example, completely different annotation methods have been shown to introduce an abundance of ‘orphan’ genes in post-hoc integrated pangene sets (Weisman et al. 2022). However, it is not clear whether annotation methods that are nominally similar (e.g., all being “integrative” approaches that combine multiple evidence types) will produce similar false positive results, particularly within species where genomic sequences are of high quality and per base sequence diversity is low. Understanding these effects is critical for accurately interpreting pangenome analyses, as methodological inconsistencies could easily be misinterpreted as biologically meaningful variation among closely related genomes.

Combined, these results demonstrate the importance of gene family structure in the evolution of even closely related plant genomes. Given the clear predisposition of plant genomes to evolve PAV (and the likely evolutionary importance of this PAV), we opted to focus on angiosperm pangenomes as a case study for the relative impacts of methodology on inference of gene PAV.

### Cotton and soybean as model systems for studying annotation-derived PAV

To test the relative impacts of methods and sequence evolution, we compared gene presence-absence variation in two aggregated sets of integrative annotations: soybean (*n* = 37, Supplementary Table 1) and cotton (*n* genomes = 11, Supplementary Table 2). Like sets of reference genomes for most species, these resources were not built in one coordinated effort. Instead, these cotton and soybean genomes were respectively built and published by several groups of scientists, each using their own methodologies (hereon ‘consortia’). This provides an excellent opportunity to examine how different methodologies affect the inference of pangenome variability. Moreover, these crops are ideal for our analysis because their limited genetic diversity (due to domestication bottlenecks) provides a controlled background against which methodological effects can be more clearly distinguished from biological variation.

Despite being constructed using different sequencing approaches and computational methods, the genome assemblies in these datasets are highly contiguous (Supplementary Table 1-2). Each consortium also generated integrated annotations by incorporating the three major lines of evidence (homology, *ab initio* prediction, and transcriptome data), with all cotton and two soybean genomes having native RNA-seq from the identical genotype used for assembly. Importantly, each individual annotation is of high quality based on standard gene prediction evaluation metrics measuring completeness (BUSCO cotton/soybean: mean = 98.6/96.7%, SD = 1.23/1.7%) and precision (PSAURON cotton/soybean: mean = 94.4/93.4%, SD = 1.4/0.74, Supplementary Table 1-2; statistics exclude 2018 Zh13) with the exception of Zh13 (Shen et al. 2018), which is an older annotation with conspicuously low BUSCO scores. This combination of high-quality assemblies with independently-generated annotations makes these datasets ideal for distinguishing methodological from biological sources of gene PAV.

### Annotation artifacts dominate biological phylogenetic signals in pangenome gene analyses

Genome-wide gene PAV should largely mirror phylogenetic relationships inferred from single copy DNA or protein sequence alignments, despite very different mutational dynamics. For example, soybean germplasm exhibits strong population structure, and Liu et al. (2020) specifically sampled a wide range of diverse ‘landrace’ locally adapted cultivars. As expected, sequence-based k-mer distances clearly reveal the expected evolutionary relationships among soybean genomes, with distinct clustering into three main groups reflecting the clear separation of cultivars based on their known genetic relationships (Fig. 1A). Furthermore, this method clearly groups the replicated nearly identical ‘Wm82’ (*n* = 3, black horizontal/vertical lines) and ‘Zh13’ (*n* = 2, white lines) assemblies.

**Figure 1.**
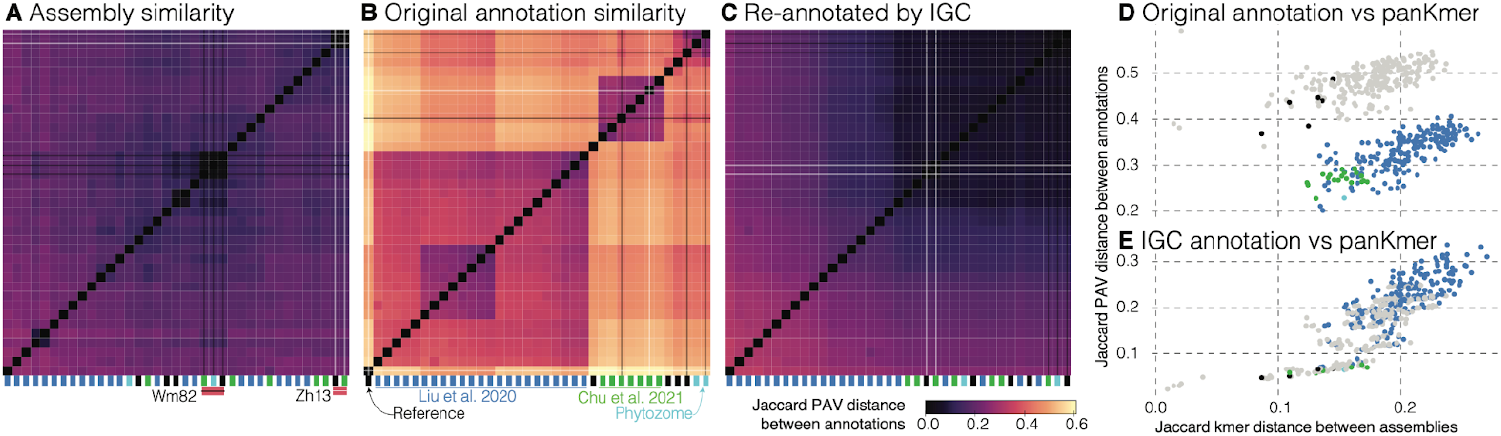
Clustering by annotation and assembly similarity reveals that original gene PAV is driven almost exclusively by annotation method. Distance matrices were calculated from panKmer (**A**) and OrthoFinder gene presence-absence variation of the originally published (**B**) and IGC (**C**) protein-coding gene annotations. In each panel, distances are hierarchically clustered to group the most similar genomes together. The annotation consortium is flagged below each column in the matrices. As a proof of concept, we analyzed redundancy among the pangenome members, which contain three versions of ‘Wm82’ (black lines) and two of Zh13 (white lines). Clear clustering of these nearly identical sequences in k-mer space, but not in annotation PAV similarity confirms the major role that gene annotation method plays when inferring biological roles of gene PAV. Correlations between the panKmer Jaccard distance and the original raw (**D**) and single-method IGC (**E**) PAV distances are presented where points are colored following panel B for comparisons within or between different consortia (grey).

However, a fundamentally different pattern emerged when we constructed trees based on gene presence-absence variation (PAV) using existing annotations hosted on SoyBase (Grant et al. 2010). Rather than clustering by evolutionary relatedness, genomes clustered into four groups almost exclusively defined by which consortium produced the annotation (Fig. 1B). This methodological signal was so strong that it entirely obscured the true biological relationships among the soybean genomes. Furthermore, the Wm82 and Zh13 assemblies annotated by different groups were more distant from each other than those from different genotypes annotated by the same consortium. We observed similar patterns in cotton genomes downloaded from Cottongen (Yu et al. 2021), though the effect was less pronounced because the particular cotton cultivars included in our study show less clear separation based on sequence and we do not have replication of assemblies with different annotation methods (Supplementary Fig. 1).

Given that annotation methods strongly influenced PAV-based clusters, it is likely that differences between the methods themselves are the driving factors. To test this hypothesis, we re-annotated all genomes using the US Department of Energy Joint Genome Institute’s (‘JGI’) Integrated Gene Calling (‘IGC’) pipeline. The resulting PAV-based trees successfully recovered the known evolutionary relationships, now closely mirroring the sequence-based trees (Fig. 1C). Critically, while assembly-based k-mer distance is completely unpredictive of PAV distance from naive integration of the original annotations (Fig. 1D; one-sided Mantel *r* = -0.12, *P* = 0.926), sequence divergence was a strong predictor of PAV from re-annotation by a consistent method (Fig. 1E; one-sided Mantel *r* = 0.861, *P* < 0.001). This result demonstrates that consistent annotation methodology is essential for accurately capturing biological signals in pangenome analyses.

### Variation in consistency among annotation methods is a major driver of estimates of pangenome ‘openness’

Biological pangenomes (i.e., all of the unique sequence present in a species) are typically classified as either ‘open’ or ‘closed’ based on how gene or sequence content changes as additional genomes are analyzed (Brockhurst et al. 2019). In a closed pangenome, the number of new genes discovered approaches zero as more genomes are added, suggesting a finite gene repertoire. In contrast, an open pangenome continues to yield new genes with each additional genome, suggesting an unbounded gene repertoire. This classification has implications for understanding species’ evolutionary potential, genomic flexibility, and breeding strategies, as it influences estimates of untapped genetic diversity and the likelihood of discovering novel beneficial alleles.

Building on our finding that annotation methodology overwhelms biological signals in phylogenetic analyses, we investigated how annotation inconsistencies affect gene PAV estimates among genomes. The genetic bottleneck followed by rare introgressions from related species in both cotton and soybean (Sreedasyam et al. 2024; Espina et al. 2024), provides an ideal setting to analyze the openness of a pangenome: while most sequence should be highly similar, low-frequency highly diverged sequences associated with adaptive introgressions are crucial targets for breeders. Furthermore, the genetic diversity present in Liu et al. (2020)’s wild and landrace genotypes should provide more opportunity to explore low frequency gene PAV in soybean. Estimates of pangenome sequence expansion from k-mers were consistent with this situation: initially the pangenomes expanded by 48/66Mb (3.4/9.6%) per cotton/soybean genome, whereas, after ten genomes were added, expansion was limited to 8.3/12.7Mb (0.5/1.4%, Fig. 2A) per genome on average. Combined, these pangenomes are not completely closed, but each genome after the first ten appear to add proportionally very few new alleles.

**Figure 2.**
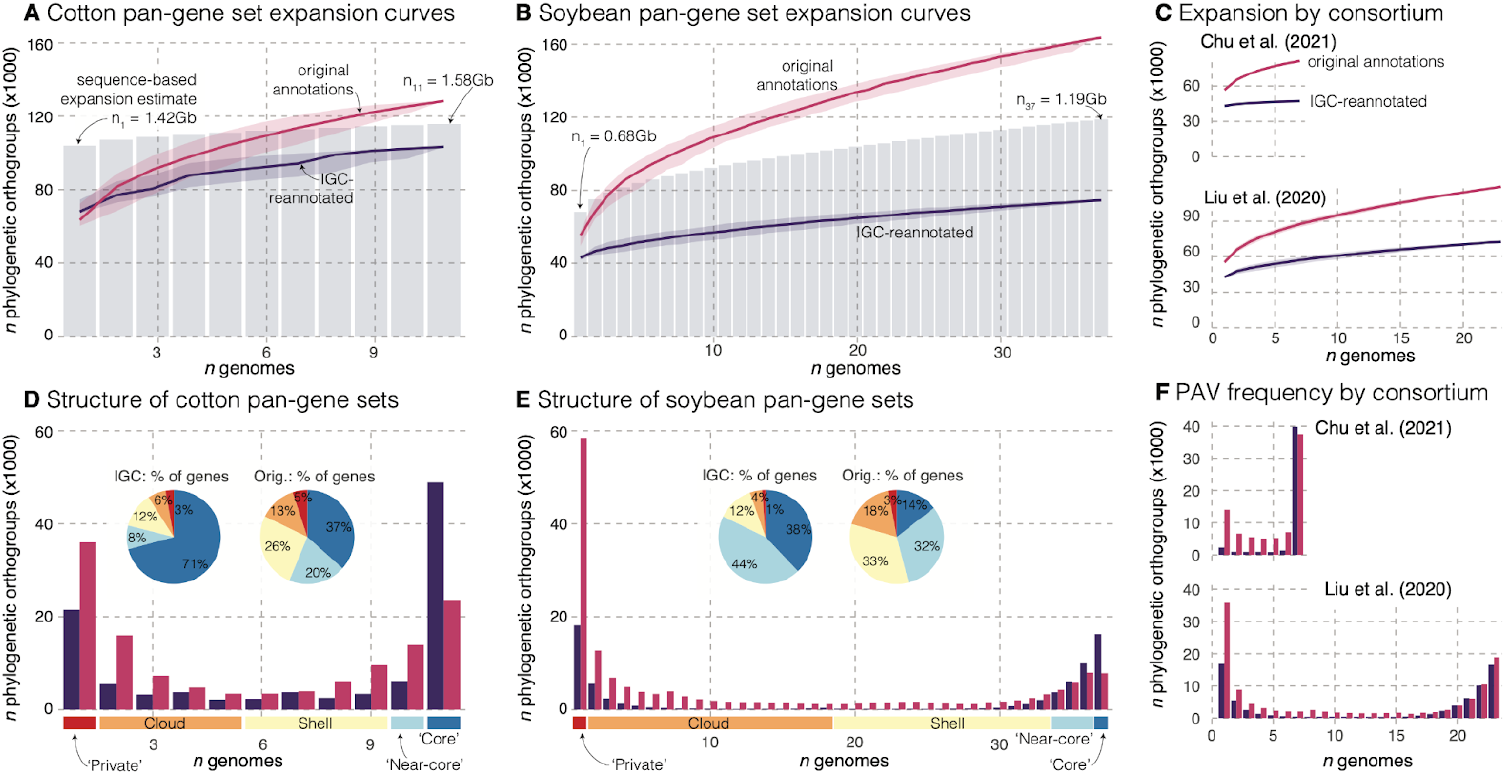
Pangenome expansion and structure in 11 cotton and 37 soybean genomes. **A** Cotton pangene set expansion curves, which depict the number of orthogroups for a given number of genomes, are presented for the original (red) and IGC-re-annotated (blue) gene sets. Solid lines and transparent polygons show mean and 95% confidence intervals of 500 simulations. The grey bar chart gives a baseline for pangenome expansion based on patterns of shared sequences in the pangenome graph and is standardized so that the total graph sequence length is the midpoint between original and IGC curve maxima (see labels for the amount of sequence represented by the smallest and largest bars). **B** and **C** follow A but for the 37 soybean genomes and annotations, and for the seven and 23 annotations by Chu et al. (2020) and Liu et al. (2021) respectively. **D** Cotton pangenome composition as a function of orthogroup presence-absence variation, categorized as private (found in a single genome), cloud (<50%), shell (<90%), near-core (≥ 90%), and core (found in all genomes) categories for both original (red) and IGC-re-annotated (blue) gene sets. Inset pie charts show the proportion of genes in each category by annotation type. **E** follow panels D but for the 37 soybean genomes and annotations. **F** follows E but for the seven genomes published in Chu et al. (2021) and the 23 in Liu et al. (2020).

In contrast to a closed sequence-based pangenome estimate, when examining pangenome growth using existing annotations, both cotton and soybean pangenomes appeared completely ‘open’, with seemingly unlimited expansion potential. For example, when using OrthoFinder’s (Emms and Kelly 2019) phylogenetically hierarchical orthogroups (HOGs), each cotton and soybean genome on average contributed 6,190 (7.8% of the average number of genes in an annotation) and 6,630 (11.8%) new HOGs respectively (Fig. 2B).

However, consistent re-annotation with a single approach dramatically transformed this picture: after annotation with the JGI IGC pipeline, each additional cotton genome contributed only 539 unique proteins (0.74% of average), while each soybean genome added just 356 proteins (0.77% of average) —reductions of ∼91% and ∼95% respectively in the rate of pangenome expansion relative to the original annotations. This IGC-reannotated pattern of pangenome expansion much more closely mirrors that of sequence-based estimates (grey bar charts in the background, Fig. 2A, B). Combined, it is clear that expansion statistics, including the ‘openness’ of a species pangenome, follow a similar pattern as genetic distance estimates: naïve integration across gene families calculated from original gene annotations produces pangenome statistics that are wildly different from sequence based estimates. Such artifacts are mostly resolved by using a consistent annotation method.

Given the massive differences between single- and multi-method annotation integration results (Fig. 1B-C), we assumed that the bulk of artifactual pangenome expansion would be due to differences among annotation methods. Indeed, each annotation method does produce many HOGs that are only found in that one consortium. For example, among the 81,104 HOGs in the Chu et al. (2021) annotations, 15,727 (19.4%) are only ever found in those seven genomes and never in the other 30 annotations. However, this was not the only driver for seemingly open cotton and soybean pangenomes. The rate of pangenome expansion *within* the seven Chu et al. (2021) and 23 Liu et al. (2020) genomes was over 6.3X and 2.1X faster in the original annotations than the IGC re-annotations respectively (Fig. 2C). Importantly, a portion of the expansion disparity between the two consortia also appears to be driven by biology: in the IGC re-annotated genomes, each Liu et al. (2020) genome contributed 1357 new HOGs, over double that of Chu et al. (2021)’s 677. This increased expansion rate may be due to the inclusion of diverse germplasm in Liu et al. (2020) but not Chu et al. (2021).

### Relative contributions of PAV categories across annotation methods

The structure of a pangenome is also often summarized by the degree and frequency of presence-absence variation (PAV), where ‘core’ sequences, which are found in all genomes, are compared to those that are in ‘private’ (found in a single genome), ‘cloud’ (< 50% presence), ‘shell’ (50-90% presence) and ‘near-core’ (>90% presence, missing in at least one genome) PAV categories. Like above, we compared a baseline expectation for the amount of sequence in PAV categories using kmer frequencies from the reference genome assemblies to PAV categories from raw annotations and re-annotated genomes with IGC.

As expected from pangenome relatedness measures (Fig. 1B), PAV calculated from integration of raw gene annotations showed high levels of presence-absence variation (Fig. 2D-E) including obvious ‘blumps’ when switching between annotation consortia (Supplementary Fig. 2). The contribution to private gene groupings was particularly conspicuous: the number of private soybean genes was nearly three times higher in the original than IGC annotations. Thus, it is not surprising that PAV as a function of membership in HOGs, was much more biased towards low frequency orthogroups in the raw than IGC annotations (Fig. 2D-E). For example, it was 3.56x (Fisher’s exact *P* < 0.001) more likely to observe low-frequency PAV orthogroups (not core or near-core) in raw than IGC soybean annotations. Importantly, ‘core’ genes were ∼2x more common in both species’ IGC re-annotated genomes than the originals. Thus, connecting orthologs and exploring candidate genes will be much more straightforward among these consistent annotations than the originals.

PAV among the IGC re-annotated genomes also more readily recapitulates biological expectations. Given sampling of more diverse material, we expected that low frequency HOGs would be much more common in the Liu et al. (2020) set than Chu et al. (2021); indeed, this pattern was clear in the IGC re-annotated genomes where 41.0% of the Liu et al. (2020) HOGs were at <50%, 4.6X higher than the 8.9% observed among Chu et al. (2021) genomes. However, these values were much closer in the original Liu et al. (2020) and Chu et al. (2021) annotations where 55.6% and 33.2% of genes had > 50% absences respectively. Combined, the relatively enriched amount of high-frequency sequences in IGC annotations meshes far more closely with evolutionary expectations and sequence-level differences, indicating that both the apparent openness of pangenomes and the extent of PAV can be substantially influenced by annotation methodology.

### Quantifying the consistency of annotations

Our analyses of phylogenetic trees and pangenome composition strongly suggest that existing annotations contain high levels of artifactual variation. In many cases, the cause is clear. For example, sequence variation can lead to minor differences in annotation heuristics support scores causing stochasticity in borderline-quality genes presence-absence (Lovell et al. 2021, 2018). However, in genomes like cotton with very little sequence variation, this artifact alone cannot explain the 6,000+ private genes observed in many genomes.

To better understand the proximate causes of gene annotation inconsistency, we examined loci with identical coding sequences between pairs of genomes and measured their predicted coordinate consistency and overlap (see Methods). We expected nearly all genomes and gene models to be consistent, given that these genes represent evolutionarily conserved, identical sequences. However, this was clearly not the case. In all comparisons, at least 13k genes mapped perfectly (with 31k on average), ensuring that our consistency analyses were based on a large subset of the gene space; however, on average across all cotton genome pairs, only 44.5% of genes with identical DNA sequence had identical structure to the original gene predictions; HPF17-Bar32 was the extreme case where just 30.8% of identical sequences had identical gene model structure (Fig. 3A). Much of this inconsistency was attributable to differences in gene annotation methods, and consistency improved markedly with our IGC cotton annotations (from 44.5 to 87.0% agreement on average, Fig. 3A-B), although some discrepancies persisted. We observed similar patterns in soybean genomes (Fig. 3C-D) where the percentage of perfectly mapped genes with identical prediction in the target genome was 61.5% for original annotations versus 96.5% in our re-annotated set. While such structural differences may not affect PAV estimates, even small differences in predicted exon-intron structure can significantly impact downstream analyses of gene function, expression, and evolution.

**Figure 3.**
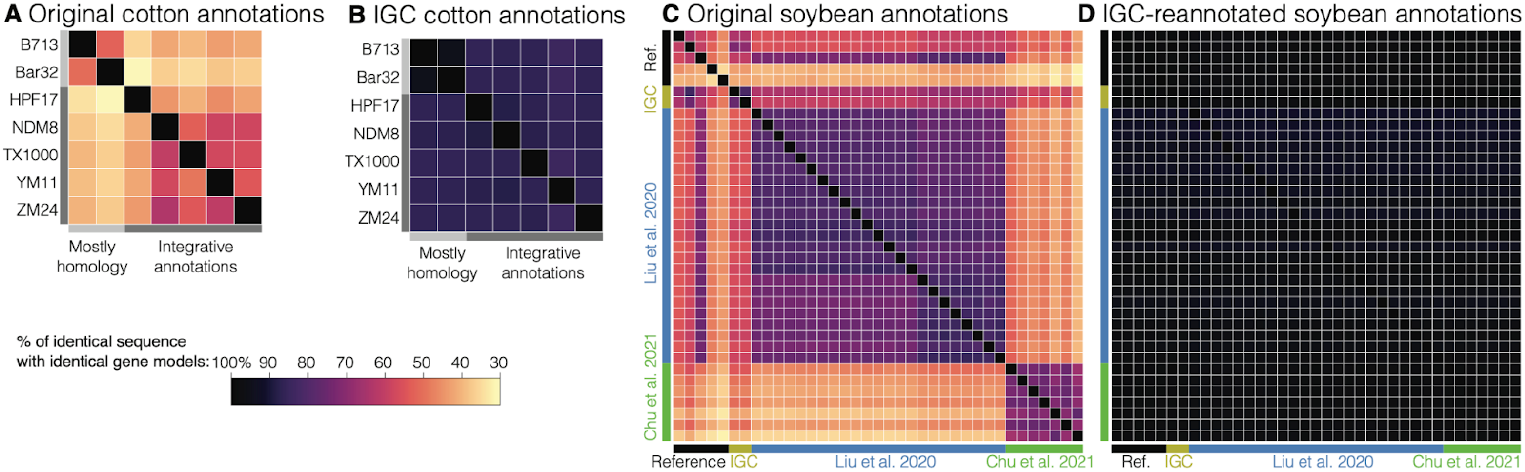
Consistency of annotations. **A** Gene models from each of the seven non-IGC cotton annotations (query, rows) were aligned to each other (target, columns) and the percent of the genes with exactly identical sequence that produced identical gene structures are reported. **B** Shows the same statistic but for the external genomes re-annotated with the IGC pipelines. **C** and **D** follow A and B respectively, except for all 37 of the soybean annotations, grouped by the consortium that produced the annotations (or ‘references’ which were built independently).

It is notable that this trend persisted even when using the more relaxed overlap metric. Instead of requiring identical gene structures, we checked whether an identical coding DNA sequence produced any overlapping coordinates with a predicted gene in the target genome on the same strand. As expected under this relaxed criterion, agreement of raw annotations increased significantly in both cotton (44.5 → 80.9%) and soybean (61.5 → 89.3%). However, the IGC-reannotated consistency of cotton (87.0 → 97.7%) and soybean (96.5 → 99.1%) increased at a corresponding level (Supplementary Figure 3).

While consistent annotation methodology dramatically reduced artifactual PAV, we also investigated whether using native RNA-seq (RNA extracted from the same genotype being annotated) affects annotation consistency. In controlled experiments with soybean YuDouNo_22, the annotation based on native RNA-seq, as opposed to RNA-seq from the reference cultivar Wm82, showed improved BUSCO completeness and PSAURON scores (Supplementary Table 3) but did not significantly affect the estimated PAV (Supplementary Figure 4). This indicates that the source of RNA-seq data is not the major driver of the reduction in PAV we observed with consistent re-annotation. Supporting this conclusion, our cotton re-annotations, which all used native RNA-seq, showed similar magnitudes of PAV reduction compared to soybean, where native RNA-seq was not always available. These findings further emphasize that methodological consistency, rather than input data sources alone, is the primary factor in reducing artifactual PAV.

## Conclusions

When examining cotton and soybean genomes, trees built from gene annotations primarily reflect who performed the annotation rather than true evolutionary relationships (Fig. 1). Annotation methodology completely overwhelms biological signal, and these methodological differences dramatically inflate estimates of gene content variation, creating another significant artifact in pangenome analysis. This finding has profound implications for comparative genomics: many studies draw evolutionary conclusions from gene content comparisons across genomes annotated by different groups, potentially leading to flawed interpretations about gene family evolution, adaptation, and species relationships. Our results highlight the critical importance of standardized annotation approaches when conducting multi-genome comparisons and challenge the reliability of conclusions drawn from heterogeneously annotated genome collections.

## Methods

### Assembly-based assessments of pangenome openness and similarity

To assess assembly-based pangenome openness, we used Pangrowth (v1.0.0) (Parmigiani et al. 2024) which calculates an exact growth curve based on a PAV matrix for all 31-mers in the input assemblies. To estimate similarity among assemblies in both sets, we used PanKmer (v0.20.4) (Aylward et al. 2023) to calculate an adjacency matrix based on the number of shared 31-mers between all pairs of genomes. We converted these adjacency values to Jaccard similarity values for assembly clustering and heatmap generation.

### Annotation-based assessments of presence-absence variation, similarity, and openness

Original (a.k.a., ‘raw’) annotations were downloaded from public repositories including Soybase (Grant et al. 2010) and Cottongen (Yu et al. 2021) and NCBI’s genome portal. The JGI’s Integrated Gene Calling pipeline (‘IGC’, see below for details) was applied to all genomes. For both the ‘raw’ and IGC annotations, a single ‘primary’ transcript was selected for each gene model, defined as the transcript with the longest coding sequence. Gene presence-absence and copy-number variation was determined by membership in OrthoFinder (Emms and Kelly 2019) defined Phylogenetically Hierarchical Orthogroups (a.k.a., ‘HOGs’), implemented through GENESPACE (Lovell et al. 2022).

### Consistent re-annotation with JGI’s Integrated Gene Calling pipeline

Seven cotton genomes were annotated with JGI’s Integrated Gene Calling pipeline (IGC) following the same protocol described in Parris et al. (2025), to match the four cotton cultivars that were originally annotated using this pipeline (FM958, TM1, UA48, and U1 HAP1; Supplementary Table 2). For each cotton genome, we used the same RNA-Seq dataset that was employed in its original annotation (Supplementary Table 1). All soybean genomes were annotated using the same protocol, but each employed a common transcriptome dataset generated for the reference Wm82.a6.v1 genome (Espina et al. 2024).

### Assessing annotation consistency

To evaluate annotation consistency between two genomes (Query and Target), we aligned the predicted coding sequences (CDS) from the Query genome to the Target genome using Liftoff (Shumate and Salzberg 2021). A CDS was considered *perfectly mapped* if it aligned with 100% sequence identity over its full length, with no gaps or mismatches. The resulting mapped gene structures were compared to the Target genome annotation using gffcompare (Pertea and Pertea 2020) in the strict mode. We calculated two metrics: *consistency*, defined as the proportion of *perfectly mapped* CDS whose genomic coordinates either exactly matched or were fully contained within those of a single predicted gene in the native annotation of the Target genome on the same strand; and *overlap*, defined as the proportion of perfectly mapped CDS that overlapped by at least one base pair with any predicted gene in the native annotation of the Target genome on the same strand, regardless of coordinate match or containment.

## Acknowledgements

The work conducted by the U.S. Department of Energy Joint Genome Institute (https://ror.org/04xm1d337), a DOE Office of Science User Facility, is supported by the Office of Science of the U.S. Department of Energy operated under Contract No. DE-AC02-05CH11231. Cotton work conducted by J.T.L. and A.S. was supported by funding from USDA-NIFA-AFRI grant (1028379) and Cotton Incorporated grant (18-753). Genome annotations for table 1 were downloaded from the following: *A thaliana*: https://ftp.ncbi.nlm.nih.gov/genomes/all/GCF/000/001/735/GCF_000001735.4_TAIR10.1/, Chicken: https://ftp.ncbi.nlm.nih.gov/genomes/all/GCF/016/699/485/GCF_016699485.2_bGalGal1.mat.broiler.GRCg7b/, Ros: https://ftp.ncbi.nlm.nih.gov/genomes/all/GCF/014/441/545/GCF_014441545.1_ROS_Cfam_1.0/, Tasha: https://ftp.ncbi.nlm.nih.gov/genomes/all/GCF/000/002/285/GCF_000002285.5_Dog10K_Boxer_Tasha/, Mouse: https://ftp.ncbi.nlm.nih.gov/genomes/all/GCF/000/001/635/GCF_000001635.27_GRCm39/, B73: https://download.maizegdb.org/Zm-B73-REFERENCE-NAM-5.0/, B97: https://download.maizegdb.org/Zm-B97-REFERENCE-NAM-1.0/, Cotton: https://ftp.ncbi.nlm.nih.gov/genomes/all/GCF/007/990/345/GCF_007990345.1_Gossypium_hirsutum_v2.1/, Coffee: https://ftp.ncbi.nlm.nih.gov/genomes/all/GCF/036/785/885/GCF_036785885.1_Coffea_Arabica_ET-39_HiFi/, Rice: https://ftp.ncbi.nlm.nih.gov/genomes/all/GCF/034/140/825/GCF_034140825.1_ASM3414082v1/.

## Description of Supplementary Material

Supplementary Tables 1-3 and Supplementary Figures 1-4 can be found in the associated online supporting material document.

## Data availability

All analysis scripts are available at https://github.com/tomasbruna/pangene-pav-integration, which retrieves all required data from Zenodo (https://doi.org/10.5281/zenodo.16809323).

