## Supplemental Material for "Evolutionary and methodological considerations when interpreting gene presence-absence variation in pangenomes"

**Supplementary Figures** **2**
Supplementary Figure 1 2
Supplementary Figure 2 3
Supplementary Figure 3 4
Supplementary Figure 4 5
**Supplementary Tables** **6**
Supplementary Table 1 6
Supplementary Table 2 7
Supplementary Table 3 7
**Supplementary References** **7**

### Supplementary Figures

#### Supplementary Figure 1

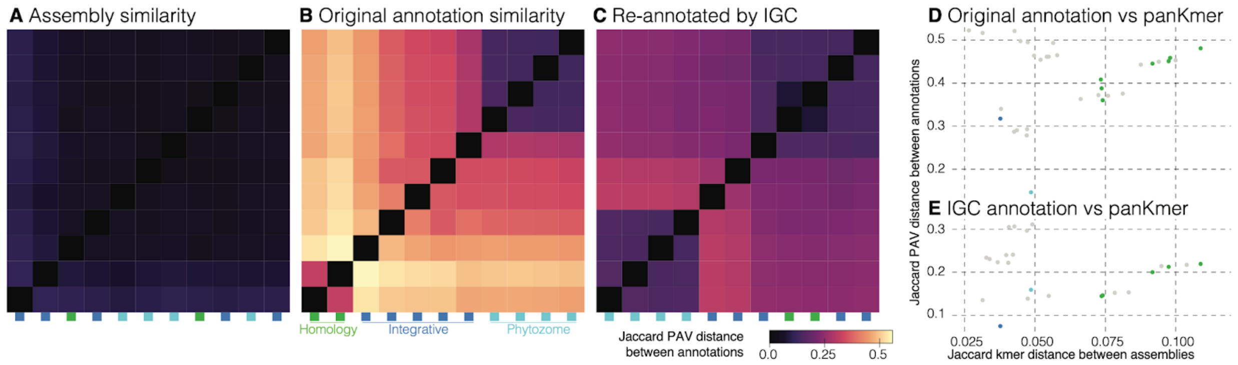


**Supplementary Figure 1 | Cotton clustering by annotation and assembly similarity reveals that original gene PAV is driven almost exclusively by annotation method.** All plots follow that of Figure 1, except with cotton data and no replication of genotypes. Distance matrices were calculated from panKmer (A) and OrthoFinder gene presence-absence variation of the originally published (B) and IGC (C) protein-coding gene annotations. In each panel, distances are hierarchically clustered to group the most similar genomes together. The annotation consortium is flagged below each column in the matrices. Correlations between the panKmer Jaccard distance and the original raw (D) and single-method IGC (E) PAV distances are presented where points are colored following panel B for comparisons within or between different consortia (grey).

#### Supplementary Figure 2

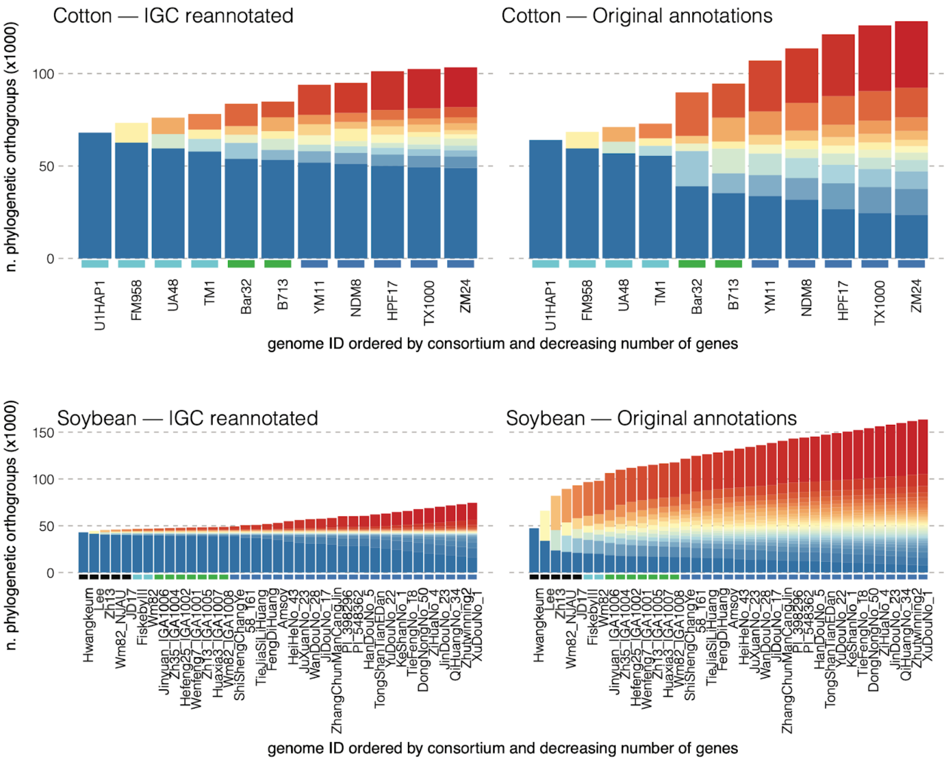


**Supplementary Figure 2 | Pangenome expansion curves grouped by consortium.** This plot follows Fig. 2A-B, but with the annotations grouped by consortium, then by decreasing numbers of annotated genes within consortium. Consortia are labeled above the gene IDs. Colors follow the pies in Fig. 2.

#### Supplementary Figure 3

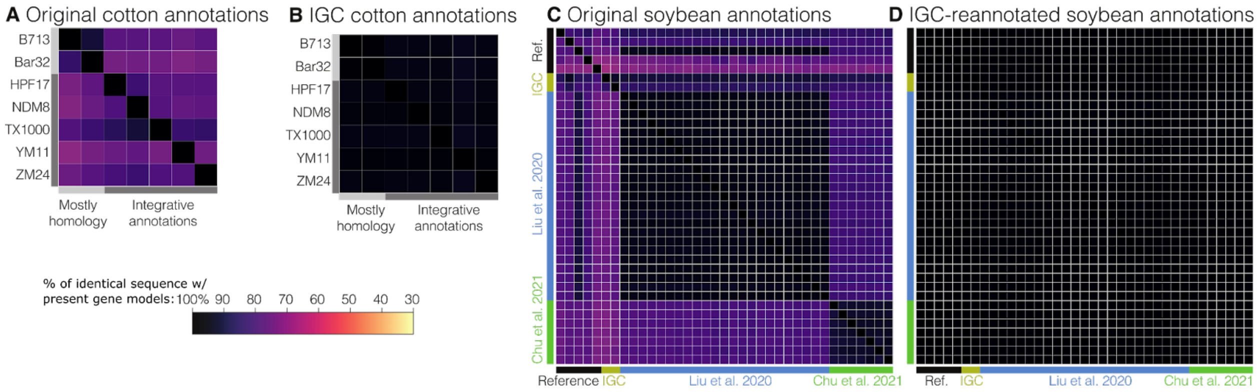


**Supplementary Figure 3 | Consistency of annotations using the relaxed overlap metric.** Consistency of annotations using the relaxed overlap metric. This figure is the counterpart to Figure 3 in the main text, but uses the relaxed “overlap” measure (see Methods).

#### Supplementary Figure 4

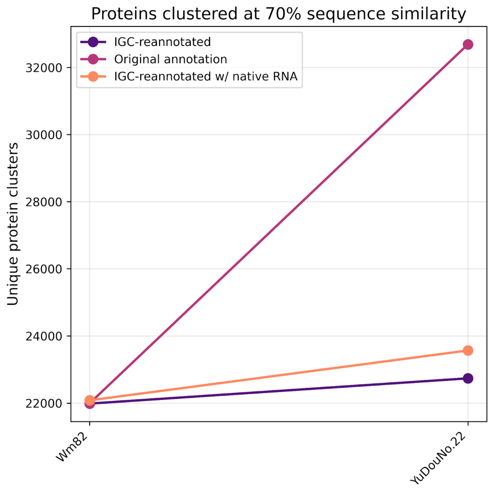


**Supplementary Figure 4 | Effect of transcriptome choice in IGC annotation on proteins newly added to the pangenome set.** Number of newly added proteins (<70% sequence similarity) to the Wm82 reference set after adding proteins predicted in YuDouNo_22 using: (a) IGC annotations guided by the Wm82 transcriptome, (b) IGC annotations guided by the transcriptome used to generate the existing annotation from Liu et al. (2020), and (c) existing Liu et al. (2020) annotation. Proteins were clustered using DIAMOND DeepClust (Buchfink et al. 2023).

### Supplementary Tables

#### Supplementary Table 1

**Supplementary Table 1 | Metadata for 37 soybean genomes.** Source information from soybase and assembly/annotation statistics calculated from the raw genomes are presented.

| Genotype ID | Consortium | Source | Assembly Size (Mb) | Assembly contig N50 (Mb) | Origi. n. genes | Orig. BUSCO | Orig PSAURON | IGC n. genes | IGC BUSCO | IGC PSAURON |
|---|---|---|---|---|---|---|---|---|---|---|
| JD17 | ref | Jidou 17 is an Chinese elite cultivar, widely grown in central China (in maturity group III). This variety is recognized for high yields and tolerance for high planting densities. | 995.3 | 18 | 52840 | 98.5 | 94.9 | 46101 | 99.6 | 95.9 |
| Wm82_NJAU | ref | Williams 82 sequenced for T2T assembly of Wang, Zhang et al. 2023 | 1,011.8 | 51.2 | 55497 | 99.4 | 90.5 | 45966 | 99.5 | 95.9 |
| Zh13 | ref | Zhonghuang 13 is a Chinese cultivar derived from accessions Yudou 18 and Zhongzuo 90052-76 by pedigree selection for high yield and stress tolerance (Shen et al., 2018; https://doi.org/10.1007/s11427-018-9360-0). | 1,011.8 | 18 | 55573 | 81.5 | 76.8 | 46292 | 99.4 | 95.7 |
| Hwangkeum | ref | Hwangkeum is an important South Korean cultivar, released in 1979 and widely used since then as a breeding parent in Korea. Hwangkeum has a determinate growth habit and nonshattering pods and is adapted to the middle Korean peninsula (Maturity Group V). | 933.1 | 7.8 | 58570 | 96.8 | 94.7 | 45766 | 99.3 | 95.9 |
| Lee | ref | Cultivar Lee, which derives from a cross of Chinese lines CNS and S-100, has been widely used as a parent in many breeding projects in the Southern U.S. and in Brazil (Wysmierski and Vello, 2013; http://doi.org/10.1590/S1415-47572013005000041). The variety is notable for resistance to Phytophthora rot, Peanut Mottle Virus, and bacterial pustule (Wysmierski and Vello, 2013). | 1,016.4 | 32.2 | 56725 | 96.6 | 93.4 | 45864 | 99.4 | 95.9 |
| Wm82 | Phytozome | Williams 82 was derived from backcrossing a phytophthora root rot resistance locus from the donor parent Kingwa into the recurrent parent Williams. A sub-line, Wm82-ISU-01, was derived by Robert Stupar (University of Minnesota) by inbreeding original W82 seed. That sub-line, registered as PI 704477, was used for assembly Wm82.gnm6 / Wm82.a6. | 1,011.1 | 44.4 | 48387 | 99.5 | 94.6 | 45952 | 99.5 | 95.9 |
| FiskebyIII | Phytozome | Bred by the late Dr. Sven Holmberg in Fiskeby, Sweden. Highly nutritious ,Äî up to 40% protein, high in calcium, iron, and vitamins (particularly A, B1, B12, and C). Thrives in northern climates. 75-80 days. | 992.2 | 15.7 | 52783 | 99.7 | 93.3 | 46024 | 99.5 | 95.9 |
| XuDouNo_1 | Liu2020 | Cultivar Xu Dou No.1 (SoyC01 from China, Jiang Su, from Liu et al. 2020 | 1,003.8 | 23.5 | 54405 | 92.4 | 93.4 | 44445 | 91.1 | 95.0 |
| Zhutwinning2 | Liu2020 | Landrace Zhutwinning2 (SoyL01 from China, from Liu et al. 2020 | 999.2 | 23.3 | 54502 | 93.0 | 93.4 | 44549 | 91.3 | 95.0 |
| QiHuangNo_34 | Liu2020 | Cultivar Qi Huang No.34 (SoyC08 from China, Shan Dong, from Liu et al. 2020 | 1,002.3 | 22.4 | 54747 | 93.8 | 93.4 | 44932 | 92.9 | 95.3 |
| JinDouNo_23 | Liu2020 | Cultivar Jin Dou No.23 (SoyC07 from China, Shan Xi, from Liu et al. 2020 | 1,008.8 | 21.8 | 54792 | 94.2 | 93.3 | 44937 | 93.7 | 95.2 |
| ZiHuaNo_4 | Liu2020 | Landrace Zi Hua No.4 (SoyL02 from China, Hei Long Jiang, from Liu et al. 2020 | 1,011.7 | 23.1 | 54803 | 94.2 | 93.4 | 45060 | 93.9 | 95.2 |
| DongNongNo_50 | Liu2020 | Cultivar Dong Nong No.50 (SoyC12 from China, Hei Long Jiang, from Liu et al. 2020 | 1,025.1 | 20 | 55000 | 94.7 | 93.5 | 45136 | 94.4 | 95.2 |
| TieFengNo_18 | Liu2020 | Cultivar Tie Feng No.18 (SoyC02 from China, Hei Long Jiang, from Liu et al. 2020 | 1,011.1 | 23.8 | 55191 | 95.6 | 93.5 | 45513 | 96.1 | 95.3 |
| KeShanNo_1 | Liu2020 | Cultivar Ke Shan No.1 (SoyC14 from China, Hei Long Jiang, from Liu et al. 2020 | 1,007.4 | 23 | 55267 | 94.9 | 93.4 | 45613 | 94.5 | 95.2 |
| YuDouNo_22 | Liu2020 | Cultivar Yu Dou No.22 (SoyC06 from China, He Nan, from Liu et al. 2020 | 1,007.9 | 23 | 55619 | 96.3 | 93.6 | 45712 | 96.6 | 95.5 |
| TongShanTianEDan | Liu2020 | Landrace Tong Shan Tian E Dan (SoyL03 from China, Jiang Su, from Liu et al. 2020 | 1,039.5 | 21.3 | 55769 | 95.5 | 93.5 | 45878 | 95.2 | 95.4 |
| HanDouNo_5 | Liu2020 | Cultivar Han Dou No.5 (SoyC09 from China, He Bei, from Liu et al. 2020 | 1,004.1 | 22.6 | 55926 | 98.3 | 93.7 | 46043 | 99.5 | 95.8 |
| PI_548362 | Liu2020 | Cultivar PI 548362 (SoyC10 from United States, Illinois, from Liu et al. 2020 | 1,004.9 | 19.8 | 56011 | 98.0 | 93.4 | 46123 | 99.5 | 95.7 |
| PI_398296 | Liu2020 | Landrace PI 398296 (SoyL05 from South Korea, Kyonggi, from Liu et al. 2020 | 1,059.8 | 22.1 | 56086 | 95.6 | 93.4 | 46038 | 95.4 | 95.4 |
| ZhangChunManCangJin | Liu2020 | Landrace Chang Chun Man Cang Jin (SoyL06 from China, Ji Lin, from Liu et al. 2020 | 997.8 | 21.5 | 56573 | 97.4 | 93.1 | 45766 | 98.0 | 95.7 |
| JiDouNo_17 | Liu2020 | Cultivar Ji Dou No.17 (SoyC11 from China, He Bei, from Liu et al. 2020 | 1,024.5 | 18.8 | 56750 | 97.8 | 93.7 | 46612 | 99.2 | 95.7 |
| WanDouNo_28 | Liu2020 | Cultivar Wan Dou No.28 (SoyC04 from China, An Hui, from Liu et al. 2020 | 1,001.5 | 22.7 | 57777 | 96.8 | 93.8 | 45692 | 98.0 | 95.7 |
| JuXuanNo_23 | Liu2020 | Cultivar Ju Xuan No.23 (SoyC03 from China, Shan Dong, from Liu et al. 2020 | 1,007.8 | 23.9 | 57792 | 96.2 | 93.7 | 45562 | 97.6 | 95.7 |
| HeiHeNo_43 | Liu2020 | Cultivar Hei He No.43 (SoyC13 from China, Hei Long Jiang, from Liu et al. 2020 | 1,010.1 | 23.8 | 57976 | 95.6 | 93.5 | 45820 | 96.6 | 95.4 |
| Amsoy | Liu2020 | Cultivar Amsoy (SoyC05 from United States, Iowa, from Liu et al. 2020 | 992.3 | 22.7 | 58040 | 96.8 | 93.7 | 45849 | 97.6 | 95.7 |
| FengDiHuang | Liu2020 | Landrace Feng Di Huang (SoyL07 from China, Ji Lin, from Liu et al. 2020 | 1,004.7 | 22.9 | 58259 | 97.1 | 93.7 | 45858 | 98.4 | 95.7 |
| TieJiaSiLiHuang | Liu2020 | Landrace Tie Jia Si Li Huang (SoyL08 from China, Ji Lin, from Liu et al. 2020 | 999.7 | 22.6 | 58496 | 97.9 | 93.6 | 46137 | 99.4 | 95.7 |
| 58_161 | Liu2020 | Landrace 58-161 (SoyL04 from China, Jiang Su, from Liu et al. 2020 | 1,005.2 | 21.8 | 58881 | 96.7 | 93.7 | 45776 | 97.8 | 95.6 |
| ShiShengChangYe | Liu2020 | Landrace Shi Sheng Chang Ye (SoyL09 from Japan, Hokkaido, from Liu et al. 2020 | 1,028.2 | 23.1 | 59588 | 98.3 | 93.7 | 47290 | 99.5 | 95.7 |
| Wm82_IGA1008 | Chu2021 | Williams 82, the soybean cultivar used to produce the reference genome sequence, was derived from backcrossing a phytophthora root rot resistance locus from the donor parent Kingwa into the recurrent parent Williams. This line, IGA1008, was sequenced and annotated by Chu, et al. (2021). | 993.0 | 1.9 | 57286 | 98.1 | 92.5 | 46221 | 99.5 | 95.8 |
| Huaxia3_IGA1007 | Chu2021 | Glycine max Huaxia 3 (IGA1007) was sequenced and annotated by Chu, et al. (2021). | 986.0 | 6.2 | 57393 | 97.5 | 92.5 | 46094 | 99.5 | 95.7 |
| Zh13_IGA1005 | Chu2021 | Zhonghuang 13 is a Chinese cultivar derived from accessions Yudou 18 and Zhongzuo 90052-76 by pedigree selection for high yield and stress tolerance (Shen et al., 2018; https://doi.org/10.1007/s11427-018-9360-0). This line, IGA1005, was sequenced and annotated by Chu, et al. (2021). | 988.8 | 4.7 | 57474 | 97.5 | 92.6 | 46085 | 99.4 | 95.8 |
| Wenfeng7_IGA1001 | Chu2021 | Glycine max Wenfeng 7 (IGA1001) was sequenced and annotated by Chu, et al. (2021). | 996.7 | 1.7 | 57505 | 97.4 | 92.8 | 46205 | 99.4 | 95.8 |
| Hefeng25_IGA1002 | Chu2021 | Glycine max Hefeng 25 (IGA1002) was sequenced and annotated by Chu, et al. (2021). | 987.3 | 2.9 | 58102 | 97.3 | 92.6 | 46205 | 99.5 | 95.8 |
| Zh35_IGA1004 | Chu2021 | Glycine max Zhonghuang 35 (IGA1004) was sequenced and annotated by Chu, et al. (2021). | 1,001.3 | 1.4 | 58150 | 97.7 | 92.8 | 46613 | 99.3 | 95.8 |
| Jinyuan_IGA1006 | Chu2021 | Glycine max Jinyuan (IGA1006) was sequenced and annotated by Chu, et al. (2021). | 995.7 | 4.3 | 58392 | 96.9 | 92.4 | 46131 | 99.5 | 95.8 |

#### Supplementary Table 2

**Supplementary Table 2 | Metadata for 11 cotton genomes.** Source information from Cottongen and Phytozome and assembly/annotation statistics calculated from the raw genomes are presented.

| Genotype ID | Consortium | Source | Assembly Size (Mb) | Assembly contig N50 (Mb) | Orig. n. genes | Orig. BUSCO | Orig PSAURON | IGC n. genes | IGC BUSCO | IGC PSAURON |
|---|---|---|---|---|---|---|---|---|---|---|
| TM-1 | Phytozome | G. hirsutum TM-1, cotton genetic standard was sequenced and annotated by Sreedasyam et al., 2024. | 2,278.1 | 40 | 75,854 | 99.8 | 94.4 | 71,247 | 99.8 | 94.4 |
| FM958 | Phytozome | G. hirsutum FM958, a conventional cotton cultivar developed and released by the Commonwealth Scientific and Industrial Research Organization (CSIRO) was sequenced and annotated by HudsonAlpha Institute for Biotechnology and Cotton Inc. Reference genome and annotation is available on Phytozome. | 2,297.3 | 69.1 | 75,965 | 99.8 | 94.0 | 72,246 | 99.8 | 94.0 |
| UA48 | Phytozome | G. hirsutum UA48 released by the Arkansas Agricultural Experiment Station was sequenced and annotated by Sreedasyam et al., 2024. | 2,293.5 | 8.1 | 77,237 | 99.7 | 93.9 | 71,995 | 99.7 | 93.9 |
| U1HAP1 | Phytozome | G. hirsutum U1, a highly FOV4-resistant upland cultivar was sequenced and annotated by Parris et al., 2025. | 2,293.4 | 89 | 71,650 | 99.8 | 94.1 | 72,315 | 99.8 | 94.1 |
| ZM24 | n/a | G. hirsutum acc. ZM24 was sequenced and annotated by Yang et al., 2019. | 2,308.2 | 1.9 | 73,707 | 99.5 | 95.2 | 70,882 | 99.8 | 96.1 |
| YM11 | n/a | G. hirsutum YM11, an accession with elite cold tolerance was sequenced and annotated by Wang et al., 2024. | 2,343.1 | 88.9 | 84,720 | 98.5 | 91.0 | 76,335 | 99.7 | 95.6 |
| TX1000 | n/a | G. hirsutum race punctatum accession no. Punctatum 25 (TX-1000) was sequenced and annotated by Peng et al., 2022. | 2,292.5 | 11.4 | 74,520 | 97.7 | 95.4 | 70,997 | 99.8 | 96.0 |
| NDM8 | n/a | G. hirsutum NDM8 was sequenced and annotated by Ma et al., 2021. | 2,291.8 | 13.1 | 79,729 | 99.0 | 93.5 | 70,248 | 99.8 | 96.2 |
| HPF17 | n/a | G. purpurascens HPF17 is a primitive race of G. hirsutum was sequenced and annotated by Cheng et al., 2024 | 2,558.7 | 10.1 | 79,146 | 97.5 | 95.1 | 79,426 | 99.8 | 95.4 |
| Bar32 | USDA | G. hirsutum Bar32 is a nematode-resistant line sequenced and annotated by Perkin et al., 2021. | 2454.3 | 75.3 | 75,988 | 97.8 | 95.4 | 70,881 | 99.8 | 96.0 |
| B713 | USDA | G. hirsutum B713 is a nematode-resistant line sequenced and annotated by Perkin et al., 2021. | 2,296.1 | 68.8 | 71,531 | 96.0 | 96.3 | 70,816 | 99.8 | 96.1 |

#### Supplementary Table 3

**Supplementary Table 3 | Effect of transcriptome choice in IGC annotation on BUSCO completeness and PSAURON score.** BUSCO completeness and PSAURON scores are shown for: (a) IGC annotations guided by the Wm82 transcriptome, (b) IGC annotations guided by the transcriptome used to generate the existing annotation from Liu et al. (2020), and (c) the existing Liu et al. (2020) annotation.

| Genotype ID | Consortium | Origi. annotated genes (n) | Orig. BUSCO (%) | Orig pSAURON | IGC annotated genes (n) | IGC BUSCO (%) | IGC PSAURON | BUSCO Improvement | PSAURON improvement |
|---|---|---|---|---|---|---|---|---|---|
| YuDouNo_22: IGC w/ transcriptome from Wm82 | Liu2020 | 55,619 | 96.3 | 93.6 | 45,712 | 96.6 | 95.5 | 0.3 | 1.9 |
| YuDouNo_22: IGC w/ native transcriptome from Liu et al. (2020) | | | | | 45,835 | **97.8** | **96.4** | **1.5** | **2.8** |
